# Mucus cell and ionocyte precursors migrate between epithelial cells to disperse through the zebrafish epidermis

**DOI:** 10.64898/2026.07.22.740112

**Authors:** Khaled Y. Nassman, Olivia Justynski, Sina Huxhagen, Sukriti Kapoor, Melissa Emami, Chanyue Hu, Matteo Pellegrini, Jeffrey B. Rosa, Alvaro Sagasti

## Abstract

Mucus-secreting cells and ionocytes play critical roles in many organs. Both cell types are usually distributed as scattered, solitary cells within an epithelium, a pattern presumably optimal for their function. To investigate how they attain their dispersed distributions, we imaged mucus cell and ionocyte precursors in the epidermis of developing zebrafish. Previous reports found that precursors of both cell types are first detected in the ventral epidermis (covering the yolk) before spreading dorsally, but the mechanism driving this progression was unknown.

Photoconverting basal epidermal cells in the ventral embryo revealed that some cells actively migrate away from this area to populate the rest of the epidermis. These cells lose their basal cell identity when they start migrating and begin expressing markers of mature mucus cells or ionocytes during migration. These cells travel entirely between the two epithelial layers of the epidermis, occasionally pause migration to divide, and repel one another through contact inhibition of locomotion, behaviors that likely aid in their dispersal. After migrating for about a day, mucus cell and ionocyte precursors intercalate into the superficial epithelial layer of the epidermis, where they complete differentiation. These observations reveal how mucus cells and ionocytes achieve their scattered distributions in the zebrafish epidermis, suggesting that similar processes promote their distribution in other mucosal organs.

## Introduction

Epithelial tissues often contain a mosaic of dispersed, specialized cell types. Broadly dispersed cellular patterns form in developing tissues through diverse mechanisms: spatially distributed cellular domains can arise from interactions between activating and inhibitory signals that obey “Turing pattern” rules (Kondo and Miura, 2010; Turing, 1952), from mechanical constraints that promote emergent patterns (Shyer et al., 2013; Shyer et al., 2017), or, perhaps most directly, from active cell movement. For example, cells in developing kidney epithelia disperse over short distances through a process termed “mitosis-associated cell dispersal”, in which cells delaminate from the epithelium, divide, and reinsert into the same epithelium a few cells away from their origin (Packard et al., 2013). To disperse over larger distances, many cell types, including neural crest cells (Carmona-Fontaine et al., 2008), cajal-retzius cells (Villar-Cerviño et al., 2013), endodermal cells (LaBelle et al., 2025), and multiciliated cells in frog epidermis (Chuyen et al., 2021), couple cell migration with mutual inhibition, a strategy called “contact inhibition of locomotion” (CIL) (Stramer and Mayor, 2017).

Mucus cells and ionocytes are specialized cell types critical for the health of many organs, including lungs, kidneys, intestines, and the inner ear (Ma et al., 2018; Pou Casellas et al., 2023; Shah et al., 2022). These cells are integrated within epithelia and interact with the external environment: Mucus cells (also called goblet cells) secrete mucus, which spreads and adsorbs over the apical surfaces of neighboring epithelial cells (Ma et al., 2018), while ionocytes (also called mitochondria-rich cells or intercalated cells in kidneys) express transporters that pump ions into and out of organs to regulate their ionic composition (Shah et al., 2022).

Although both cell types are relatively rare within a tissue, mucus cells and ionocytes are crucial for organ health. For example, misregulation of mucus secretion or ion transport in lungs causes debilitating diseases, including cystic fibrosis, asthma, and COPD (Johnson, 2011; Ma et al., 2018; Montoro et al., 2018; Plasschaert et al., 2018; Yuan et al., 2023). In many organs, mucus cells and ionocytes derive from similar basal stem cell populations that express the transcription factor tp63 (Blomqvist et al., 2004; El-Dahr et al., 2017; Montoro et al., 2018; Plasschaert et al., 2018; Quigley et al., 2011), and must translocate into superficial epithelial layers to reach the external surface of the tissue. Mucus cells and ionocytes are specified by foxa (Lai et al., 2016; Swisa et al., 2024; van der Sluis et al., 2008; Wan et al., 2004; Ye and Kaestner, 2009) or foxi (Hsiao et al., 2007; Hulander et al., 2003; Jänicke et al., 2007; Pou Casellas et al., 2023; Quigley et al., 2011; Vidarsson et al., 2009) family transcription factors, respectively, and share molecular profiles across organs (Pou Casellas et al., 2023). Both cell types are often distributed broadly within epithelia as discrete cells (rather than in clusters), a pattern presumably critical for their function. How these cells attain their dispersed distributions is not known.

The epidermis in zebrafish embryos is a bilayered, mucosal epithelium, consisting of a tp63-expressing basal cell layer and a superficial periderm layer (Le Guellec et al., 2004; O’Brien et al., 2012). In larval zebrafish, mucus cells and several types of ionocytes are interspersed within the periderm cell layer in embryos and larvae. In addition to the skin, mucus cells are also found in the gastrointestinal tract and the pharynx, but each of these organs uses distinct mechanisms to specify mucus cells (Lai et al., 2016). Similarly, specialized ionocytes are also found in neuromasts (Peloggia et al., 2021; Peloggia et al., 2024) and olfactory epithelium (Peloggia et al., 2025), but, compared to epidermal ionocytes, ionocytes in those organs use different specification mechanisms, have a distinct paired organization, and derive from dedicated stem cell populations. The specification of epidermal mucus cells and ionocytes is well characterized: Notch signaling promotes mucus cell development (Lu et al., 2021) and inhibits ionocyte development (Hsiao et al., 2007; Jänicke et al., 2007), and, like in mammals, mucus cells are specified by foxa proteins (Lai et al., 2016), whereas ionocytes are specified by foxi proteins (Esaki et al., 2009; Hsiao et al., 2007; Jänicke et al., 2007). Markers for mucus cells and ionocytes first appear in the ectoderm above the yolk, then spread posteriorly past the yolk extension, before appearing more broadly through the embryo in a ventral-to-dorsal sequence (Hsiao et al., 2007; Jänicke et al., 2007; Lu et al., 2021; Solomon et al., 2003).

Ionocytes are subdivided into several functionally distinct subtypes, each of which expresses different ion transporters (Horng et al., 2007; Hwang and Chou, 2013; Liao et al., 2007; Liao et al., 2009; Lin et al., 2006; Wang et al., 2009). Each of these cell types are distributed in distinct patterns over broad regions of the epidermis. Intriguingly, specialized ionocytes in sensory neuromast organs derive from basal cells in the adjacent epidermis and migrate extensively within neuromasts before intercalating into the superficial epithelium in polarized positions (Peloggia et al., 2021; Peloggia et al., 2024), raising the possibility that ionocyte precursors in the epidermis may also migrate before integrating into the periderm.

To investigate how mucus cell and ionocyte precursors attain their broad distributions, we monitored cells derived from the basal layer of the zebrafish epidermis using time-lapse imaging. We found that mucus cell and ionocyte precursors segregate from the basal cell population, usually after cell division, and migrate for hours through the epidermis before intercalating into the periderm cell layer. These cells move with a protrusive morphology in the interepithelial space between basal and periderm cells and begin differentiating into their respective cell types before integration. These observations raise the possibility that similar migratory processes promote the distribution of mucus cells and ionocytes in mammalian organs.

## Results

### Mucus cell and ionocyte precursors arise in the ventral embryo and migrate to disperse through the epidermis

To characterize mucus cells and ionocyte distribution in the zebrafish epidermis, we first used RNA-scope fluorescent in situ hybridization to confirm and extend the reported progression of ionocyte and mucus cell marker expression. Previous work demonstrated that early markers for ionocyte specification (foxi3a and foxi3b) first appear in cells covering the yolk region during early somitogenesis stages, then spread through the epidermis in a ventral-to-dorsal sequence. Like in mammals (Rajavelu et al., 2015; van der Sluis et al., 2008; Wan et al., 2004; Ye and Kaestner, 2009), foxA family transcription factors specify mucus cells in zebrafish (Lai et al., 2016), but which of three foxA family members (foxa1, a2 or a3) is required specifically for epidermal mucus cell development is unknown. Based on RNA-seq data (see below, Figure 4), foxa1 and foxa3 were the likeliest candidate for mucus cell specification in the skin, and foxa3 was robustly expressed in the epidermis by *in situ* hybridization (Figure 1A, Supplemental Figure 1). Co-staining at later stages with known mucus cell markers (agr2 and muc5.1) confirmed that foxa3 marks epidermal mucus cell precursors (not shown). Staining for ionocyte (foxi3a and foxi3b) and mucus cell (foxa3) precursors confirmed that these two populations are distinct from each other and follow the reported developmental progression--appearing first over the yolk, then extending posteriorly past the yolk extension, before spreading dorsally (Figure 1A, Supplemental Figure 1). Intriguingly, foxa3 also appeared early in a few cells in the head (between the eyes and otic vesicle), raising the possibility that there may be a second origin for mucus cell precursors in the anterior embryo. To characterize the spread of differentiating mucus cells, we stained embryos for transcripts reported to be specific to mucus cells (agr2, (Lai et al., 2016; Lu et al., 2021)) and two types of ionocytes: atp1b1b for Na+/K+-ATPase-rich cells (NARCs) and atp6v1aa for H⁺-ATPase-Rich Cells (HRCs) (Hsiao et al., 2007; Jänicke et al., 2007; Lin et al., 2006) (Figure 1B, Supplemental Figure 2). Unexpectedly, many cells expressed both agr2 and atp1b1b or atp6v1aa. We confirmed that these cells were mucus cells, since agr2 cells expressed foxA transcription factors. We conclude that some mucus cells express these “ionocyte” genes, defining a distinct subset or developmental stage of mucus cell precursors (and suggesting that previous studies of ionocytes may have included some mucus cells).

**Figure 1.**
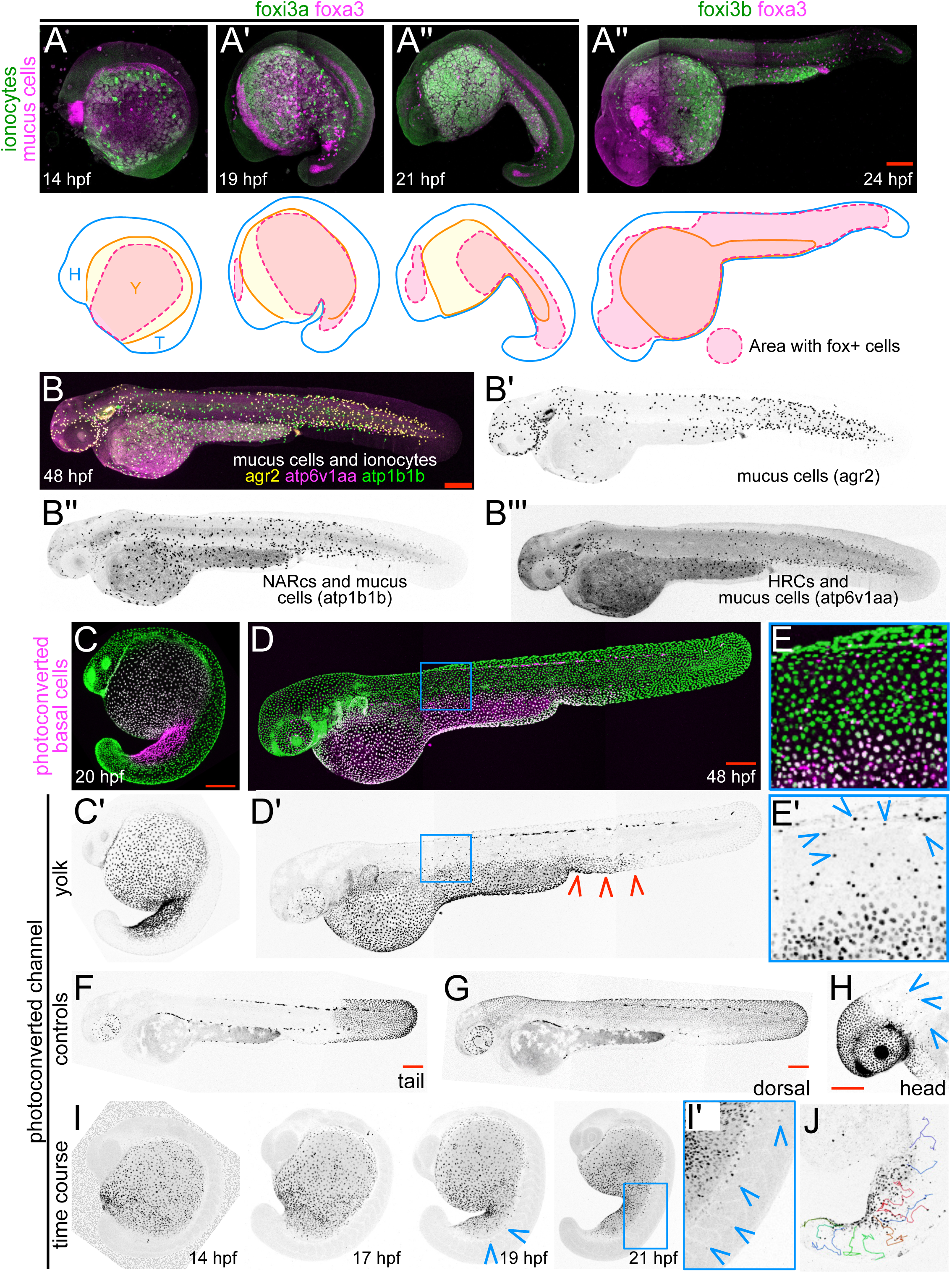
Mucus cells and ionocytes spread away from the ventral embryo. A) RNA-scope in situ hybridization for genes expressed in mucus cell (foxa3) and ionocyte (foxi3a or foxi3b) precursors at the indicated stages. Diagrams below roughly depict regions where mucus cell and/or ionocyte precursor cells are found (red shaded regions). Note that the first three timepoints use foxi3a to represent ionocyte precursors, while the last panel uses foxi3b, since foxi3a diminishes and foxi3b becomes more widespread over time. For a more comprehensive gene expression time series for all three genes, see Figure S1. B) RNA-scope in situ hybridization at 48 hpf for agr2 (B’, a mucus cell reporter), atp1b1b (B’’, expressed in NARC ionocytes and mucus cells) and atp6v1aa (B’’’, expressed in HRC ionocytes and mucus cells). Note that since all three reporters are expressed in mucus cells at this stage, each cell type can only be distinguished in the merge (B), where ionocytes are identifiable by their lack of agr2 expression. For a more comprehensive gene expression time series of all three genes, see Figure S2. C-J) Following basal cell derivatives with photoconversion. Basal cell nuclei in the yolk region (tp63[BAC]:Gal4FF;UAS:nls-Eos) were photoconverted at 20 hpf (C, C’); by 48 hpf the photoconverted cells had spread posteriorly past the yolk extension (red arrowhead) (D, D’) and individual cells had migrated dorsally away from the photoconverted region (E, E’) (e.g., blue arrowheads). E, E’ are insets of the boxed region indicated in D and D’. All basal cells are shown in C-E; the photoconverted channel alone is shown in C’-E’. F-G) Basal cell regions in the tail (F) or dorsal embryo (G) that were photoconverted at 20 hpf remained contiguous regions at 48 hpf. H) “Representative image of a fish in which basal cells were photoconverted in the head at 20 hpf, showing several photoconverted cells that had separated from the photoconverted region, indicating that cells may also migrate away from this region. I) Time course of basal cells photoconverted in the yolk region at 14 hpf shows that cells begin to migrate away from this region between 17 and 19 hpf (arrowheads). I’ is an inset of the boxed region in 21 hpf timepoint. J) Tracing of migratory cell trajectories from a time-lapse imaging of photoconverted cells (photoconversion at 18 hpf, movie starts at 20 hpf) superimposed onto the last time frame of the movie.

Nonetheless, we could distinguish each cell type by the combination of genes they expressed. Cells expressing these reporters appeared roughly in the same locations as the fox transcription factors, but appeared later in development (Figure 1B, Supplemental Figure 2). Consistent with previous reports (Hsiao et al., 2007; Jänicke et al., 2007; Kowalewski et al., 2021; Lu et al., 2021), by 48 hpf HRCs (expressing atp6v1aa only) were concentrated around the yolk, NARCs (expressing atp1b1b only) were densest in the trunk, and mucus cells (expressing agr2) were spread throughout the animal but most dense in the head and tail.

The spread of ionocyte and mucus cell precursors through the epidermis could occur via a patterned differentiation sequence and/or active migration. To distinguish these possibilities we labeled basal cell nuclei with the photoconvertible fluorescent protein Eos (tp63[BAC]:Gal4FF; UAS:nls-Eos) and photoconverted nuclei in the entire yolk region (n= 7) or just the yolk extension (n = 7) at 18-20 hpf (Figure 1C-C’). By 48 hpf, the photoconverted region had extended posteriorly, beyond the yolk extension, but some photoconverted cells were located dorsally, separate from the contiguous photoconverted region (Figure 1D-E’). By contrast, photoconverted cells in the tail or dorsal embryo did not disperse from photoconverted patches (Figure 1F-G). In the head, in 5 out of 6 experiments at most one cell had segregated away from the photoconverted patch, but in one experiment several dispersed cells were seen away from the photoconverted patch (Figure 1H), potentially reflecting a second population of epidermal mucus cells, as suggested from the foxa3 expression pattern (Supplemental Figure 1). Collectively, these results suggest that photoconverted basal cells or their descendants migrate away from the yolk extension to dorsal positions.

To determine when cells begin migrating dorsally away from the yolk, we photoconverted the yolk at an earlier stage (14 hpf), and tracked cells over the next several hours (n = 6) (Figure 1I). Photoconverted cells began appearing outside of the photoconverted region between 17 and 19 hpf (Figure 1I-I’). Time-lapse movies confirmed that individual photoconverted cells migrated away from the photoconverted tissue region (Figure 1J, Movie 1). We conclude that migratory cells emerge from basal cells and begin migrating dorsally ∼18 hpf. This temporospatial progression resembles that of cells specified to become mucus cells (foxa3-expressing) and ionocytes (foxi3a-expressing) (Figure 1A, Supplemental Figure 1).

To further map the trajectories of basal cell-derived migratory cells, we photoconverted basal cells at later stages in different parts of the embryo (Supplemental Figure 3). Photoconverting cells at the top of the head at 24 hpf and following them by time-lapse for 12 hours revealed that most migrating cells move from ventro-lateral regions towards the top of the head (n = 2, Supplemental Figure 3A-B). Photoconverting regions in the dorsal and ventral trunk at 20 hpf and following them by time-lapse for 11.5 hours revealed that more cells migrate ventral-to-dorsal than dorsal-to-ventral (n = 5 dorsal and 5 ventral photoconversion experiments, Supplemental Figure 3C-E). These observations strengthen the conclusion that basal cell-derived migratory cells travel away from ventral regions to populate the rest of the embryo (Supplemental Figure 3F).

### Migratory cells emerge after division of tp63+ cells and lose their basal cell identity

To investigate how basal epithelial cells give rise to migratory cells, we imaged basal cell nuclei (tp63[BAC]:Gal4FF;UAS:nls-GFP) in the region of the yolk extension, where our photoconversion experiments suggested many migratory cells arise. In several instances, we noted that stationary cells appearing to be basal cells divided, giving rise to two daughter cells, both of which began migrating a few hours after cell division (Figure 2A-B, Movie 2). Thus, migratory cells arise from basal cells and the onset of migration appears to be coupled to cell division.

**Figure 2.**
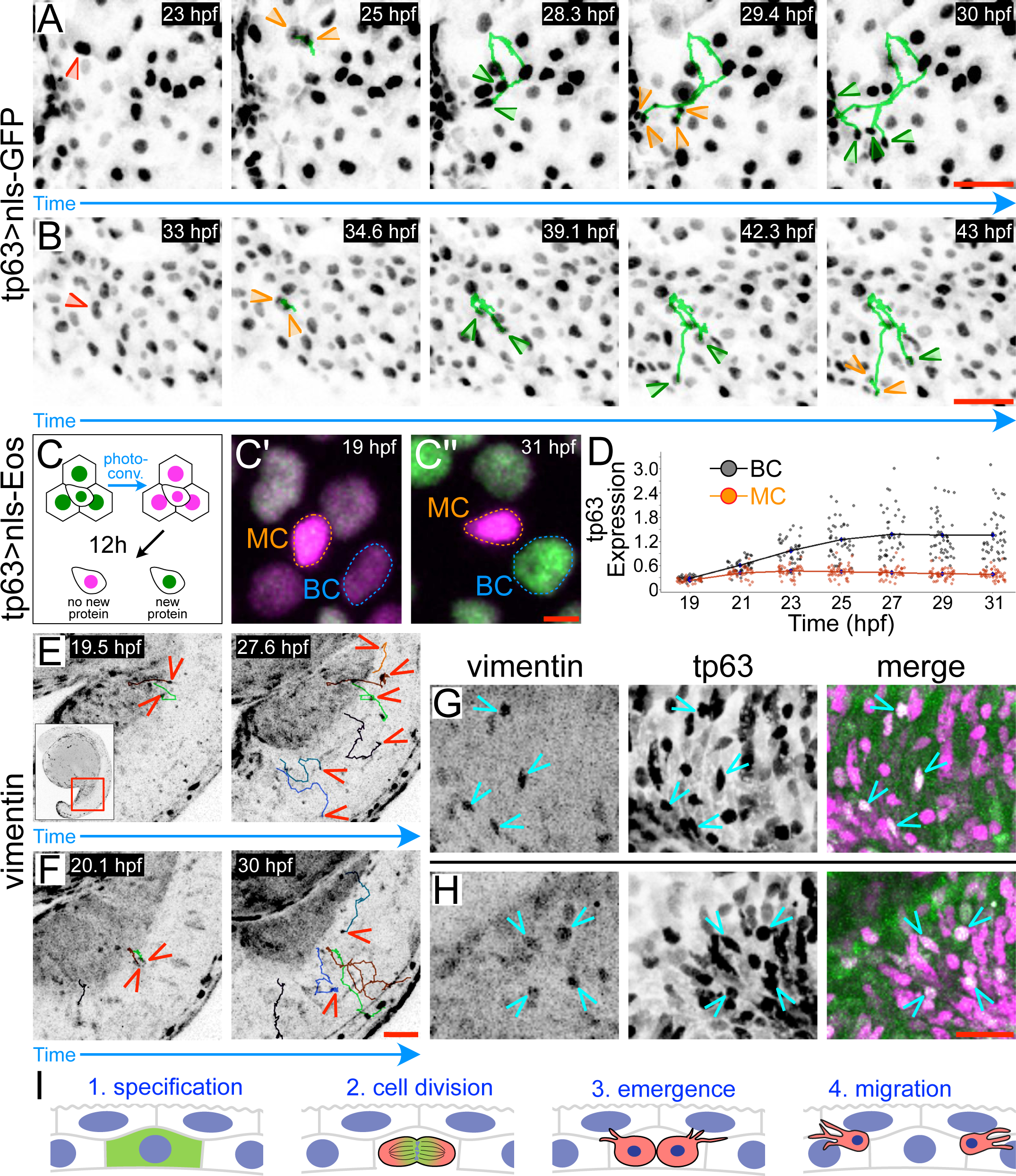
Migratory cells emerge after division of tp63+ cells and lose their basal cell identity. (A-B) Stills from two representative time-lapse movies of cells expressing a nuclear basal cell reporter (tp63[BAC]:Gal4FF; UAS:nls-GFP). Stationary cells within the basal epithelium (red arrowheads in first panels) divide (orange arrowheads in second panels), then both daughters begin migrating (green trace and arrowheads). Many migratory cells divide again, often within hours of emergence (orange arrowheads in 4th panel in A, and in 5th panel in B). Note that migratory cell emergence usually occurs before 24 hpf (A), but can sometimes occur later in development (B). C) Experimental schematic: Eos-expressing basal and migratory cells (tp63[BAC]:Gal4FF; UAS:nls-Eos) were photoconverted from green to red at 19 hpf and time-lapse imaged for 12 hours. Newly synthesized Eos fluoresces green; cells in which the tp63 reporter had turned off continued to fluoresce red but never fluoresced green again. C’-C’’) Images just after photoconversion (C’) and 12 hours later (C’’); MC=migratory cell; BC=basal epithelial cell. D) Quantification of green to red fluorescence ratio (tp63 expression) at indicated time points post-photoconversion in stationary basal cells (grey) and migrating cells (orange). E-F) Stills from two different time-lapse movies of a vimentin reporter line (-2vim:eGFP) (LeBert et al., 2018). Inset shows the entire animal; red box indicates the approximate location of main panels. Traces show migratory cell trajectories; arrowheads show location of migratory cells at the indicated time-point. G-H) Images of two different embryos expressing the vimentin reporter and tp63 reporter (tp63[BAC]:Gal4FF; UAS:nfsb-mCherry). Arrowheads indicate cells expressing both reporters. I) Model for migratory cell emergence from basal epidermal cells: (1) basal cells are specified during early somitogenesis stages; (2-4) At ∼18 hpf, these cells divide and both daughters begin migrating between basal and periderm cell layers. Scale bars in A-B, G-H = 40 μm; Scale bar in C = 5 μm; Scale bars in E-F = 50 μm.

Epithelium-derived cells that become migratory change their transcriptional program as they lose their epithelial identity (Kalluri and Weinberg, 2009). Although we discovered these migratory cells using a recombinant BAC reporter for tp63, tp63 itself maintains basal cell identity, so we suspected that the reporter may no longer be active when cells are migrating. To test this possibility, we photoconverted nuclear Eos driven by the tp63 driver (tp63[BAC]:Gal4FF;UAS:nls-Eos) at 19 hpf and time-lapse imaged embryos for 12 hours (Figure 2C-C’’). By the end of the movie, most epithelial nuclei contained newly synthesized green fluorescent Eos, but migratory cells contained only the red photoconverted Eos (Figure 2C’, C’’, D), indicating that although the reporter was still active in basal cells, it had shut down in migratory cells. We are thus able to image migratory cells with this reporter only because fluorescent protein made in precursor cells perdures.

Epithelial-to-mesenchymal transition is characterized by stereotyped changes in gene expression, including downregulation of keratin and upregulation of vimentin intermediate filaments (Kalluri and Weinberg, 2009). To test if the emergence of these migratory cells shares this characteristic, we time-lapse imaged a vimentin reporter line (LeBert et al., 2018). Several cells expressing this reporter indeed migrated dorsally away from the yolk extension (Figure 2E, F, Movie 3). Crossing vimentin and tp63 reporter lines confirmed that the two reporters are co-expressed in migrating cells (Figure 2G, H), indicating that the emergence of migratory cells from tp63-expressing basal cells shares at least one molecular characteristic with EMT and further supporting the conclusion that migratory cells emerge near the yolk extension. Together, our observations indicate that migratory cells derived from tp63-expressing basal cells change fate as they begin migrating (Figure 2I).

### Mucus cell and ionocyte precursors differentiate as the migrate

We hypothesized that basal cell-derived migratory cells are ionocyte and mucus cell precursors, given the similarity in their spatial and temporal trajectories (Figure 1). To test this hypothesis, we obtained a reporter line for HRC ionocytes (CEACAMz1:mCherry-F (Kowalewski et al., 2021)) and made a new transgenic allele with a previously described reporter transgene for mucus cells (-6agr2:GFP (Lai et al., 2016)). We crossed each of these lines to fluorescent reporters driven by the tp63 BAC driver and made time-lapse movies. Both reporters started to appear in basal cell-derived migratory cells (labeled with tp63[BAC]:Gal4FF; UAS:nls-GFP or UAS:nfsb-mCherry) during migration (Figure 3, Movie 4), several hours before they became stationary and intercalated. These observations confirm that basal cell-derived migratory cells become mucus cells and ionocytes, and indicate that cells begin differentiating as they migrate.

**Figure 3.**
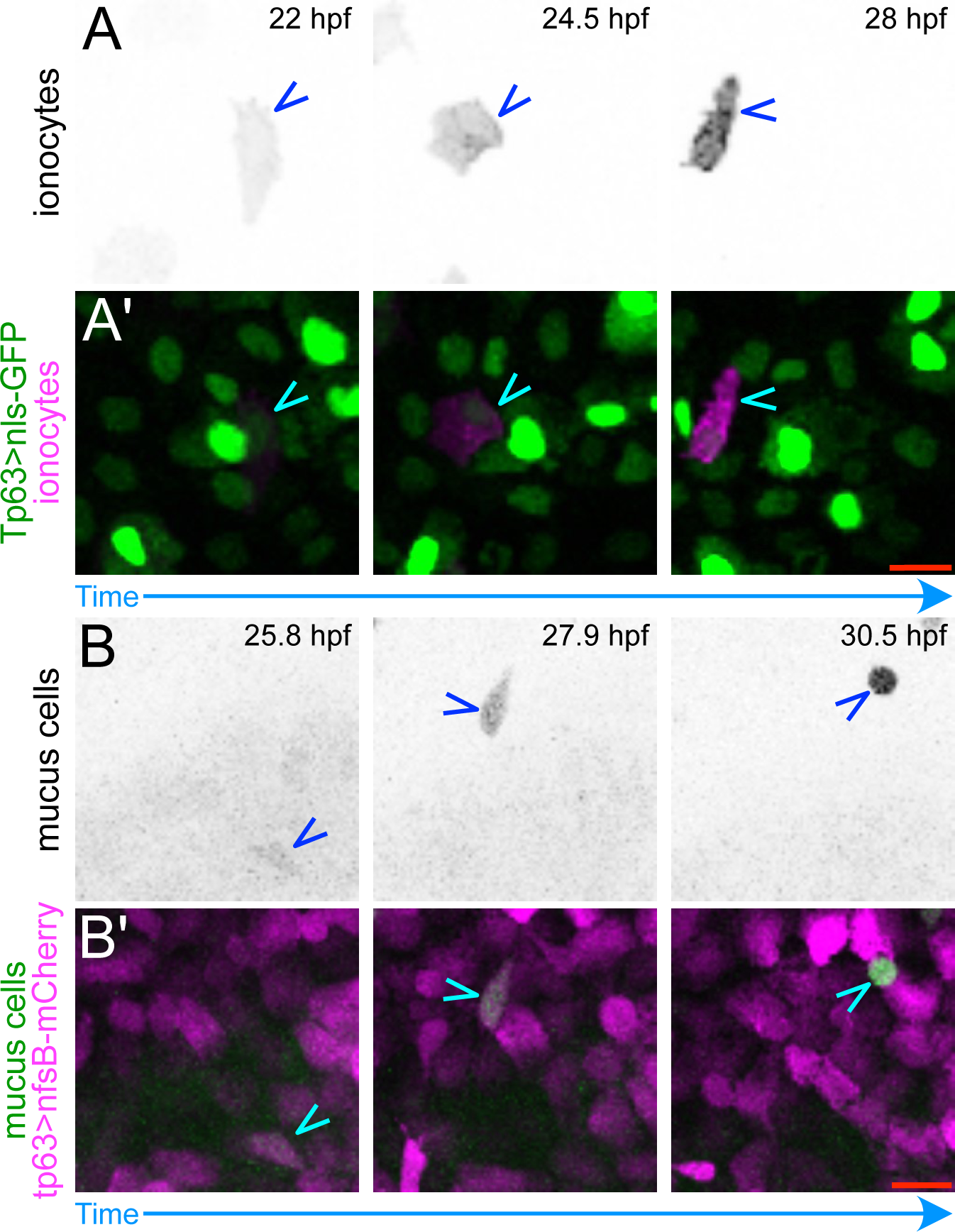
Migratory cells express reporters for ionocytes and mucus cells during migration. A) Stills from a time-lapse movie show that a reporter for HRC ionocytes (CEACAMz1:mCherry-F) (Kowalewski et al., 2021) turns on in basal-derived migratory cells (p63[BAC]:Gal4; UAS:nls-GFP) during migration. A shows ionocyte precursor channel alone; A’ shows merge. Arrowheads point to ionocyte precursor cells. B) Stills from a time-lapse movie show that a mucus cell reporter (-6agr2:GFP) (Lai et al., 2016) turns on in basal-derived migratory cells (p63[BAC]:Gal4; UAS:nfsb-mCherry) during migration. B shows mucus cell precursor channel alone; B’ shows merge. Arrowheads point to mucus cell precursor cells. Scale bars = 20 μm.

**Figure 4.**
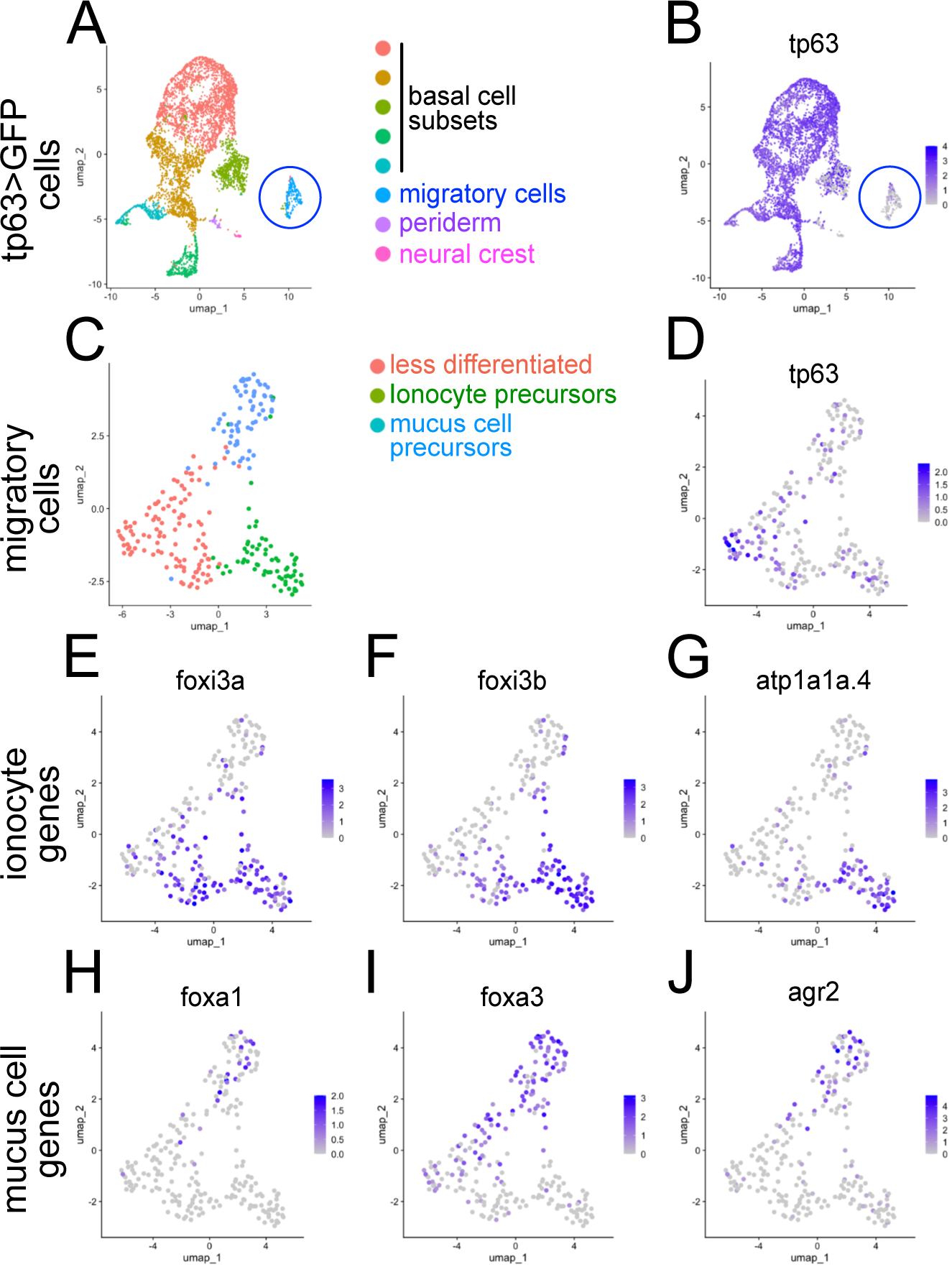
Single cell RNA-seq identifies gene expression in mucus cell and ionocyte precursors. A) UMAP plot of GFP-expressing cells purified from a basal cell reporter (tp63[BAC]:Gal4FF;UAS:nls-GFP). tp63>GFP cells segregate into 8 clusters. Five clusters appear to be regional subpopulations of ectodermal basal cells. The circled cluster contains candidate migratory mucus cell and ionocyte precursors. B) Feature plot showing tp63 expression. The cluster containing presumed migratory cells expresses relatively low levels of tp63 and is indicated with a blue circle. C) UMAP plot of migratory cells (circled cluster in A-B) subdivided into three clusters. D-J) Feature plots showing relative expression of tp63 (D), 3 ionocyte genes (E-G), and 3 mucus cell genes (H-J) in the migratory cell cluster.

To further characterize the differentiation of migrating mucus cell and ionocyte precursors, we used fluorescence-activated cell sorting to purify cells expressing a tp63-driven reporter (tp63[BAC]:Gal4FF; UAS:nls-GFP) at 24 hpf, and subjected them to single cell RNA-sequencing (Figure 4). Cells segregated into 8 clusters, most of which, as expected, robustly expressed tp63 (Figure 4A-B). Cells in five clusters expressing high levels of tp63 were contiguous; each of these clusters may contain basal cells from different parts of the animal (e.g., cells in the fin fold or sensory placodes) or in different cell cycle states. Two small clusters with low tp63 expression were likely periderm and neural crest cell contaminants, based on expression of characteristic genes. One cluster, consisting of 241 cells, also expressed relatively low levels of tp63 and was thus a candidate for the tp63-derived migratory cell population (Figure 4B, circled cluster).

Further dividing this candidate cluster into three subclusters (Figure 4C) revealed that tp63 expression was higher in one subcluster (Figure 4D), which also contained cells that expressed either the mucus cell specification gene foxa3 or the ionocyte specification gene foxi3a. Notably, although this subcluster contained both mucus cell and ionocyte specification genes, individual cells contained one or the other. This gene expression signature is consistent with this subcluster consisting of less differentiated mucus cell and ionocyte precursors. Another subcluster was enriched in the ionocyte specification gene foxi3a, as well as foxi3b and atp1a1a.4, markers of more mature ionocytes (Figure 4E-G). The third subcluster was enriched in foxa3, and some cells expressed foxa1 and/or agr2, also markers of mucus cells (Figure 4H-J). We thus conclude that this migratory cell cluster contains distinct ionocyte and mucus cell precursors at various stages of differentiation, providing a resource for investigating gene expression in these developing cells. Since mature mucus cells and ionocytes have markedly distinct functions and morphologies, the fact that their precursors cluster together indicates that they are transcriptionally similar, likely reflecting their migratory properties and shared descent from epidermal basal cells.

### Migratory cells travel between epithelial layers in eccentric paths with protrusive morphologies

To determine where within the epidermis mucus cell and ionocyte precursors migrate, we imaged basal and migratory cells expressing membrane-localized eGFP (tp63[BAC]:Gal4FF;UAS:EGFP-PLCδ-PH) and periderm cells expressing membrane-localized RFP (krt5:mp-mCherry) (Rosa et al., 2023) (Figure 5A-B). All migratory cells (>20/20) were sandwiched between the two epithelial layers of the epidermis. We never saw cells migrating along the basement membrane below basal cells, or between lateral membranes of basal or periderm cells. We next imaged mucus cell precursors specifically (-6agr2:GFP) along with basal cells (tp63[BAC]:Gal4FF;UAS:nfsb-mCherry) and rendered the images in 3D with Imaris software (Figure 5C-C’), which confirmed that these cells migrate on the apical surface of basal cells. Thus, the migration of mucus cell and ionocyte precursors takes place exclusively in the 2-dimensional intraepithelial region of the epidermis.

**Figure 5.**
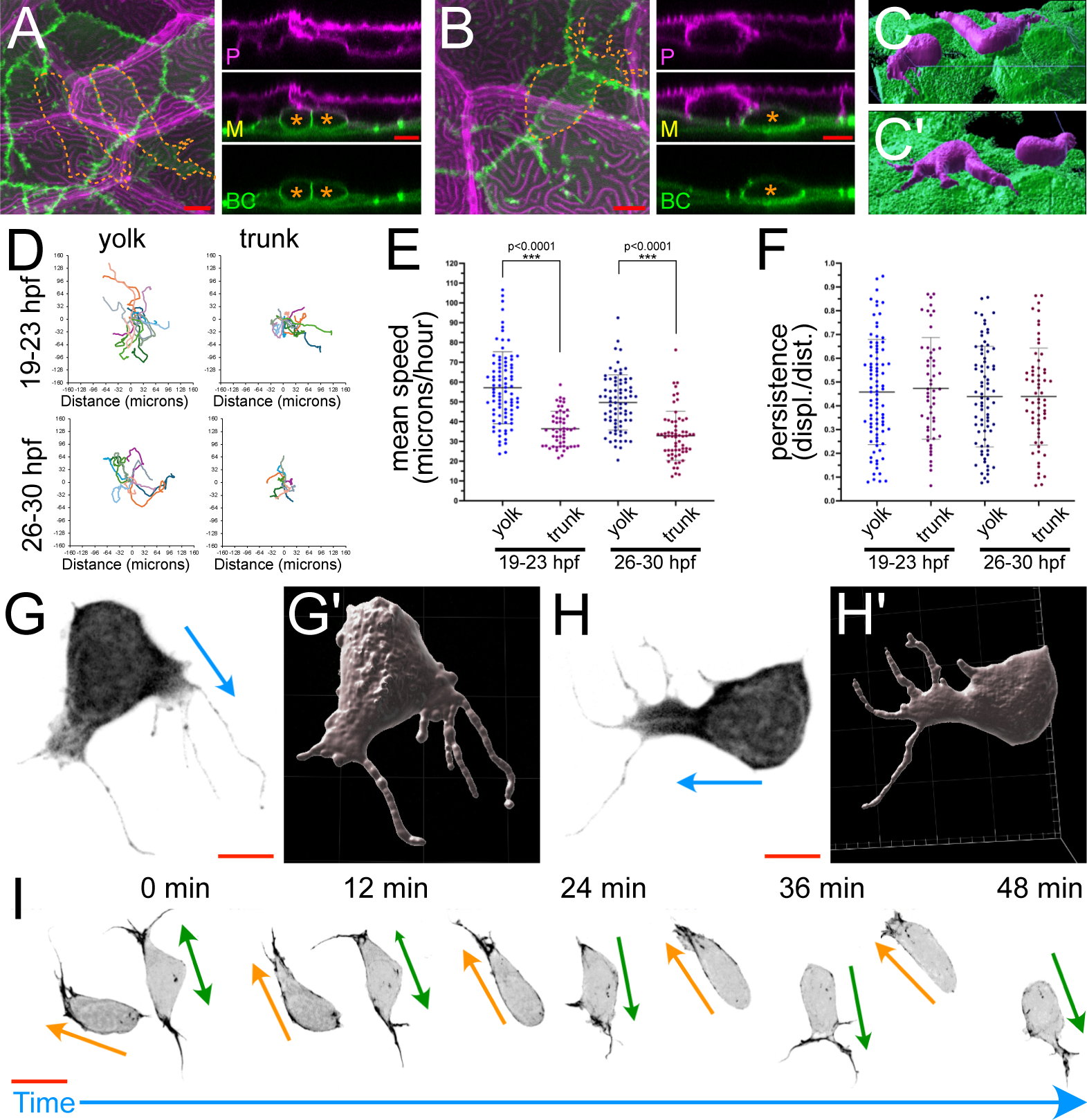
Mucus cell and ionocyte precursors migrate within the developing epidermis. A-B) Images of cells with labeled periderm (magenta) and basal and migratory cell membranes (green). *En face* view is on the left, orthogonal views on the right. P = periderm, BC = basal and migratory cells, M = merge. Asterisks indicate migratory cells. Scale bars in A and B = 10 μm. C-C’) 3D rendering of images of mucus cell precursors (magenta) with basal cells (green). D) Traced migratory cell trajectories in the yolk and trunk regions at the indicated stages. Traces are aligned at their origin point. E) Plot of mean speed at the indicated times and locations. Each dot represents mean velocity for an individual cell (n=8 embryos for yolk and 7 embryos for trunk at both timepoints). F) Plot of persistence (displacement divided by the total trajectory length) at the indicated times and locations. Cell population analyzed is the same as in D. G-H’) Migratory cell morphologies: two representative migratory cells imaged with -6agr2:GFP. Image is shown to the left (G,H), 3D reconstruction (Imaris) is shown to the right (G’, H’). Arrows indicate the direction of migration. Scale bars in G and H = 5 μm. Corresponding time-lapses are in Movie 5. I) Stills from a time-lapse movie of migratory cells with labeled actin. Arrows indicate the directions of migration. Scale bar = 10 μm.

To characterize migration, we made time-lapse movies of nuclei labeled with the tp63 driver in different parts of the animal at different stages and traced their trajectories. Overlaying migratory paths of migratory cells revealed no obvious directional bias (Figure 5D). Most cells migrated between 20 and 100 μm/hour (Figure 5E). Average migration speeds differed in different parts of the embryo, and migration velocities decreased over time (Figure 5E). Path persistence (net displacement/total path length) ranged from very straight paths (close to 1) to highly tortuous paths (close to 0), but the range of path persistence measurements was similar at different stages and in different parts of the embryo (Figure 5F).

To better appreciate the morphology of migrating mucus cell and ionocyte precursors, we imaged migrating mucus cell precursors (-6agr2:GFP) starting at ∼28 hpf, at higher resolution. Cells migrated with highly polarized morphologies, with lamellar and filopodial protrusions at their leading edges and often rounded rear membranes (Figure 5G-H) (Movie 5). 3D renderings showed that leading protrusions usually extended within the plane of the epidermis, between periderm and basal cells (Figure 5G’,H’). When migrating mucus cell precursors encountered one another, they repolarized and moved away from each other, a behavior characteristic of contact inhibition of locomotion (CIL) (Stramer and Mayor, 2017) (Movie 6). To image their actin cytoskeleton, we injected a reporter transgene (UAS:Lifeact-GFP) into fish with a basal cell driver (tp63[BAC]:Gal4FF), resulting in mosaic expression. Actin was concentrated in dynamic protrusions at the leading edge, whereas the back of cells contained little actin (Figure 5I, Movie 7). Although more observations are needed to characterize their mechanism of motility, these migratory characteristics are consistent with pseudopod-based amoeboid motility (SenGupta et al., 2021).

### Migratory mucus cell and ionocyte precursors proliferate as they migrate

Time-lapse imaging revealed that, in addition to dividing just before their transition into a migratory state, basal cell-derived migratory cells often paused migration to divide and expand the population. Most, if not all, migratory cells paused to divide within a few hours of their emergence (usually between ∼20-24 hpf) (e.g., Figure 2A-B, Movie 2), and many divided a third time over the next several hours (after 24 hpf) (Movie 8). Imaging mucus cell precursors (-6agr2:GFP) also demonstrated that these cells occasionally paused migration to divide, giving rise to two GFP-expressing daughter cells (Movie 8), confirming that proliferation during migration expands the population of mucus cells. To quantify the frequency of cell division during later stages of migration, we photoconverted individual migratory cells (tp63[BAC]:Gal4FF;UAS:nls-Eos) at 24 hpf and imaged animals again 24 hours later. In 15/26 animals, a photoconverted cell remained the only red fluorescent cell at the later time-point, indicating that it had likely not divided (we almost never saw a migratory cell die). In 11/26 animals, there were 2 photoconverted cells, indicating that a single cell division had occurred.Thus, ∼40% of migratory cells divide once after 24 hpf.

### Mucus cell and ionocyte precursors intercalate into the periderm cell layer and form tight junctions with surrounding cells

Migratory mucus cell and ionocyte precursors should eventually stop migrating and integrate into the periderm, the superficial epithelial layer of the epidermis. Time-lapse movies of nuclei of basal cell-derived migratory cells (tp63[BAC]:Gal4FF; UAS:nls-GFP), revealed that most cells stop migrating by ∼40 hpf (Figure 6A), settling in stable positions between epithelial cell nuclei. Movies of migratory cells mosaically labeled with an actin reporter revealed that migratory cells intercalate into the superficial epithelial layer by wedging an apical actin-rich protrusion between periderm cells and forming an apical actin ring that expands to make an apical surface (Figure 6B, Movie 9), similar to the process described for intercalation of multiciliated cells into the frog epidermis (Chuyen et al., 2021; Sedzinski et al., 2016; Ventura et al., 2022).

**Figure 6.**
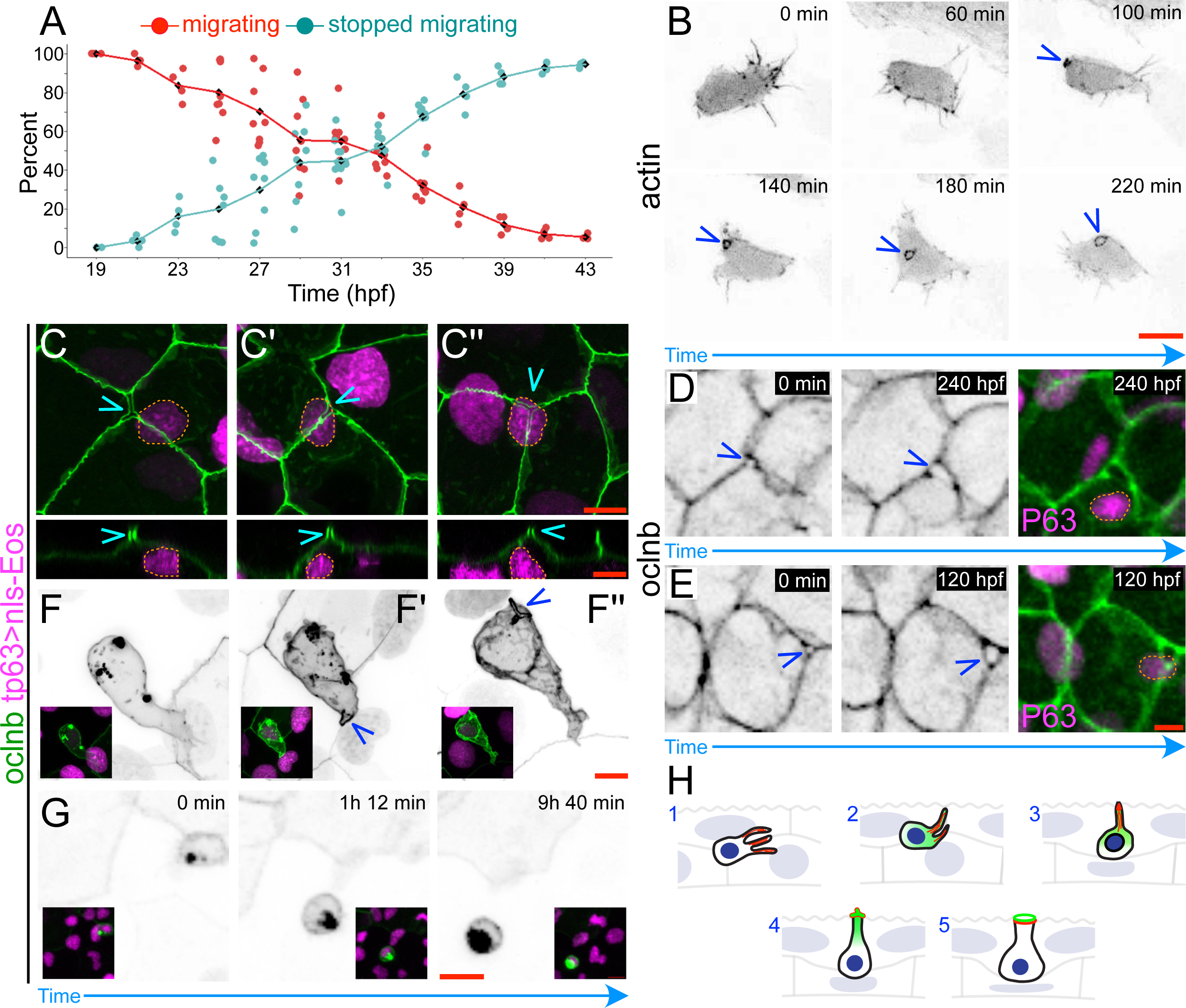
Migratory cells intercalate into the periderm cell layer. A) Proportion at indicated timepoints of migrating tp63+ cells versus formerly migrating tp63+ cells that have stopped moving. Data was quantified from time-lapse movies of the trunk (region over the yolk extension). Each data point is from different movies, lines connect mean (black diamonds) at each timepoint. B) Stills from a time-lapse movie of an actin reporter mosaically expressed in migratory cells (UAS:LifeAct-GFP transgene injected into tp63[BAC]:Gal4FF). Arrowheads indicate an expanding apical projection. C-C’’) Three examples of openings between periderm tight junctions (imaged with oclnb:GFP[BAC]) above migratory cells that have stopped moving (nuclei labeled with photoconverted tp63[BAC]:Gal4FF; UAS:nls-Eos; nuclei of formerly migratory cells are outlined). Orthogonal view is shown below. Arrowheads indicate apical openings between periderm cells. D-E) Stills from two different time-lapse movies showing that openings between tight junctions (labeled with oclnb:GFP) expand over time. Rightmost panel shows merge with labeled basal cell nuclei (photoconverted p63[BAC]:Gal4FF; UAS:nls-Eos); nuclei of formerly migratory cells are outlined. F-F’’) Migratory cells mosaically labeled with a tight junction reporter (oclnb:GFP[BAC]). F shows a cell that has not yet intercalated; F’ and F’’ show cells that have begun intercalation (arrowhead indicates apical opening). Insets show merge with labeled basal cell nuclei. G) Stills from a time-lapse movie of a mosaically labeled migratory cell that expresses oclnb (oclnb-GFP[BAC]) during migration.O clnb-GFP concentrates in an apical protrusion. Insets show merge with labeled basal cell nuclei. H) Model for intercalation: Migrating cells (1) begin expressing oclnb during migration (2); cells wedge an actin- and oclnb-rich protrusion between periderm cells (2-3); the apical tip of the protrusion separates adjacent periderm cells, which often appears as a y- or x-shaped gap between periderm cells (4); the apical surface of the cell projection forms tight junctions with surrounding periderm cells and forms an expanding ring of actin at its perimeter (5). Scale bars in B and G = 10 μm; all other scale bars = 5 μm.

Imaging a BAC reporter with a GFP-tagged tight junction protein, occludin B (Oclnb-GFP[BAC]), along with nuclei of basal cell-derived migratory cells (photoconverted tp63:Gal4FF[BAC], UAS:nls-Eos), revealed that migratory cells settle underneath small gaps between periderm cells (Figure 6C), as expected for mucus cells and ionocytes. Consistent with actin imaging, these movies demonstrated that gaps in the epithelium increased in area over time after migratory cells settled below them (Figure 6D, E). Notably, Oclnb:GFP was expressed not only in periderm cells, but also at lower levels in migratory cells themselves. To confirm this observation, we injected the occludin B reporter into embryos to attain mosaic expression, and increased the saturation level of the GFP signal. This approach confirmed that Oclnb-GFP was expressed in migratory cells and concentrated in apical protrusions (Figure 6F-G) prior to intercalation, indicating that these cells likely form tight junctions with surrounding periderm cells upon intercalation (Figure 6H).

## Materials and methods

### Fish husbandry and transgenic lines

Zebrafish (*Danio rerio*) adults were housed at 28.5°C with 13.5/10.5 h light/dark cycles. Embryos were typically raised at 28.5°C in “blue water”, containing 0.06 g/L Instant Ocean salt mix (Spectrum Brands, AA1-160P) and ∼0.05% methylene blue (Thermo Fisher Scientific, 042771.AP). In some instances, embryos were raised at 25°C to slow their development. Developmental stages were approximated in hours post-fertilization at 28.5°C, but verified and corrected according to morphological criteria (e.g., somite number).

### Transgenic lines

**Table 1:**
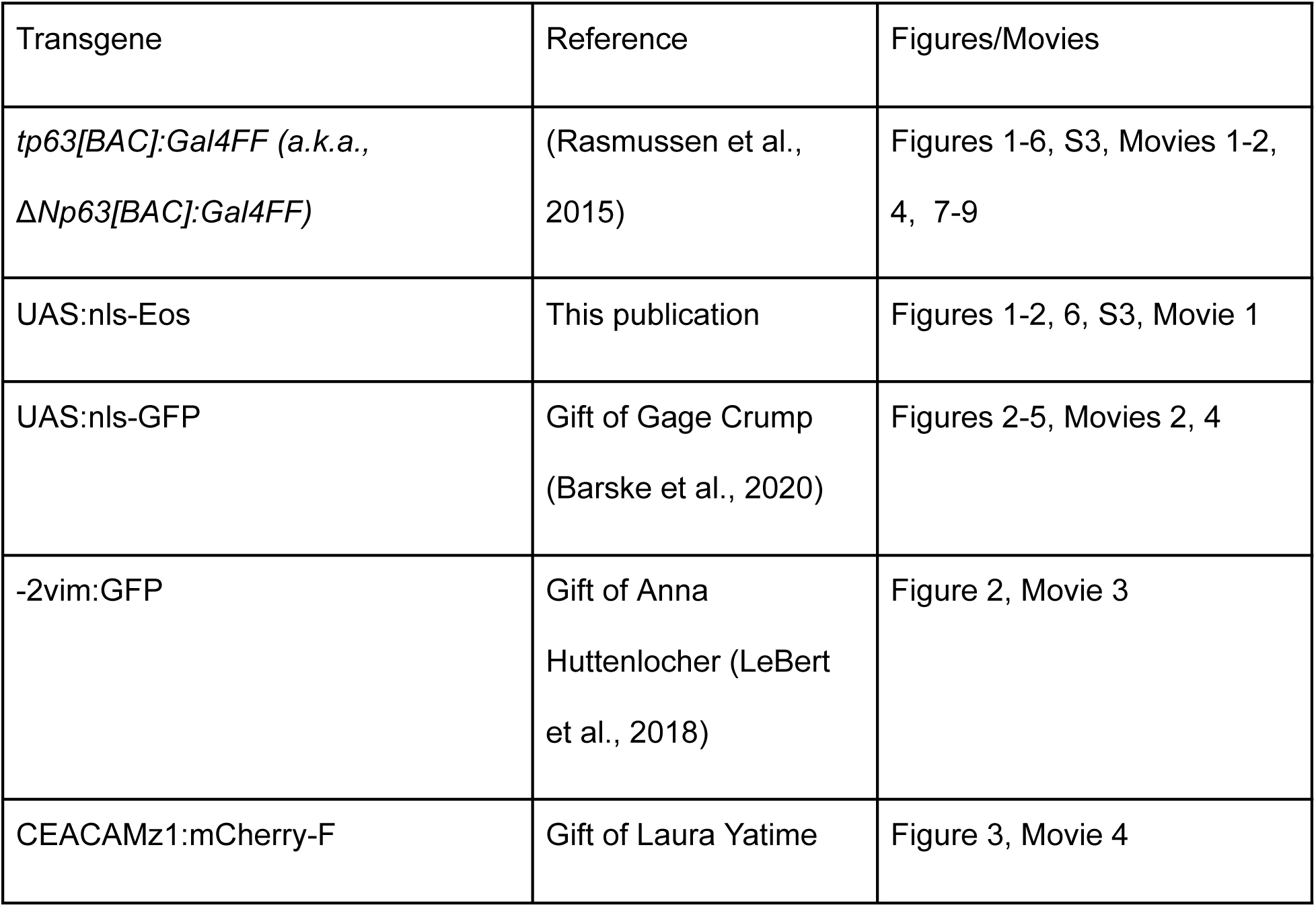
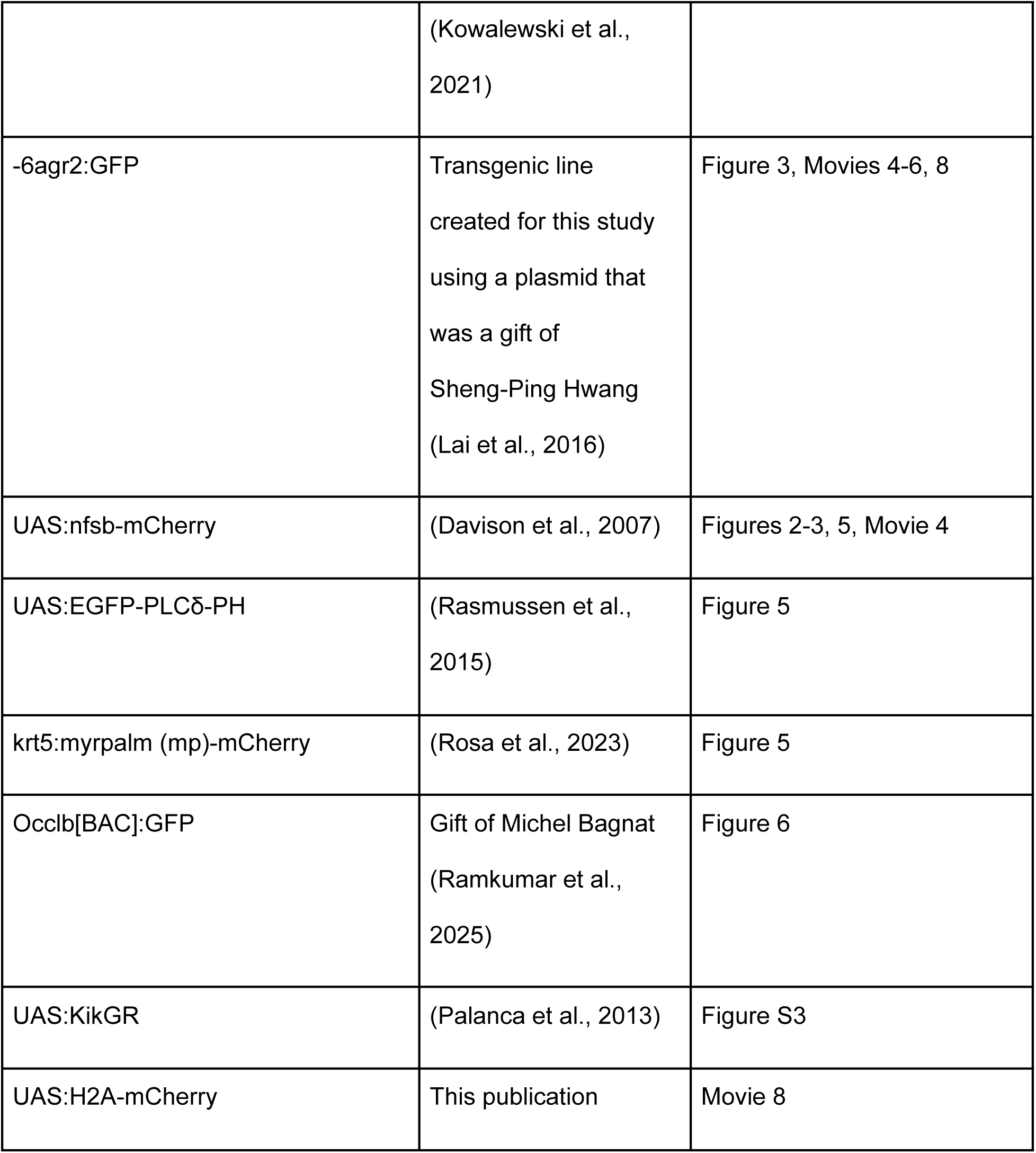
Transgenic lines used in this study.

The UAS:nls-Eos and UAS:H2A-mCherry transgenic plasmids were assembled using the tol2 Gateway system (Kwan et al., 2007). These plasmids and the -6agr2:GFP transgene (Lai et al., 2016) were injected with tol2 transposase into wildtype AB embryos to make new transgenic lines.

### RNA-scope in situ hybridization

RNAscope™ Multiplex Fluorescent V2 Assay (Advanced Cell Diagnostics) was used to detect RNA transcripts using probes targeting foxi3a, foxi3b, foxa3, atp1b1b, atp6v1aa, and agr2. Zebrafish embryos aged 14-48 hpf were fixed in 4% paraformaldehyde in PBS for 1-2 hours at room temperature, dehydrated through a graded methanol/PBS series (25%, 50%, 75%) and stored in 100% MeOH at -20°C overnight. Embryos were permeabilized with Protease Plus for 30 minutes and washed three times with 1X PBS containing 0.1% Triton X at room temperature. Next, embryos were hybridized with prewarmed 1X probes at 40°C for 2 hours. Standard RNAscope V2 multiplex reagents and 1:1500 diluted TSA Vivid Fluorophores 520, 570, and 650 were used to visualize the signal. Embryos were cleared through a graded glycerol/PBS series (25%, 50%, 75%) and mounted for imaging on a Zeiss LSM 800 confocal microscope.

### Photoconversion

Tg(tp63[BAC]:Gal4FF;UAS-nls-Eos or UAS:KikGR) embryos were anesthetized in Tricaine and mounted dorsally or laterally against a coverslip in 1.2% low-melt agarose. Immobilized embryos were placed in an imaging chamber constructed from custom-made plastic rings sealed onto coverslips with vacuum grease and filled with embryo water containing Tricaine (O’Brien et al., 2009). Regions of Interest (ROIs) were drawn with the Regions tool in the Zeiss ZEN Blue software on a Zeiss LSM 800 confocal microscope with a 10x objective. The Bleaching module was utilized to photoconvert Eos and KikGR reporters within the ROIs, using a 405 nm laser at 80% laser power for a total duration of 50 seconds. Agarose-embedded embryos were rescued using tweezers to pull apart the agarose and embryos were subsequently placed in blue water for recovery.

### Imaging

Embryos were anaesthetized in 0.4 mg/ml tricaine (MS-222, Western Chemical) and immobilized in 1-1.2% low-melt agarose in a custom-made ring chamber, sealed between a coverslip and slide with vacuum grease, as previously described (O’Brien et al., 2009). Most imaging was performed on a Zeiss LSM 800 confocal microscope at 10x, 20x, or 40x magnification. Figures 5F-G and Movies 5 and 7 were imaged on a Nikon Ax R with NSPARC at 40x or 60x magnification with 488 nm excitation laser. Movie 8 was imaged on a Zeiss LSM980. Imaris software was used to generate 3D renderings in figure 5.

### Cell sorting and RNA-seq

Embryonic zebrafish were euthanized with ice cold blue water and chorions and yolks were removed by triturating with a glass pipette. Embryos were then digested into single cells by nutating at 4°C for 20 minutes in a buffer of Dulbecco’s Modified Eagle Medium (DMEM)/F12 (Gibco, no phenol red) with 10 mg/mL Protease from *Bacillus licheniformis* (Sigma). Cells were filtered through a 40 μm cell strainer and resuspended in phosphate-buffered saline (PBS) with 2% fetal bovine serum (FBS) and 1mM EDTA for flow cytometry. DAPI (1:1000) was added immediately before sorting to exclude dead cells. Cells were sorted on a BD FACSAria with FACS DiVA software. Cells were sorted into FACS buffer and then resuspended in PBS with 0.04% bovine serum albumin (BSA) for single-cell RNA sequencing.

Single-cell RNA sequencing data from GFP+ cells was processed using the standard cellranger 9.0.1 pipeline (10X Genomics). Cells were filtered for quality control to avoid doublets and dead cells. Dimensionality reduction and downstream data visualization were completed using the Seurat package in R (Version 2025.09.2+418) (Butler et al., 2018; Hao et al., 2021; Hao et al., 2024; Satija et al., 2015; Stuart et al., 2019). Data is presented as log-normalized mRNA counts (expression).

### Cell tracking, quantification, and statistical analyses

To quantify migratory cell directionality, a region of the head in (tp63[BAC]:Gal4FF; UAS:KikGR) embryos was photoconverted at 24 hpf and imaged for 12 hours, while the dorsal and ventral halves of the trunk in (tp63[BAC]:Gal4FF;UAS-nls-Eos) embryos were photoconverted at 20 hpf and imaged for 11.5 hours. The Trackmate plugin (Ershov et al., 2022; Tinevez et al., 2017) in ImageJ/Fiji (Schindelin et al., 2012) was used to track migratory cells moving into and out of the photoconverted region, and the Cell Counter plugin (Schindelin et al., 2012) was used to quantify the number of migratory cells that had migrated along different directions at the final imaged timepoints. Data visualization was performed in R using the ggplot2 package to generate the dot plots in Figure S3B and S3E.

To quantify tp63 expression experiments, ImageJ was used to manually draw ROIs around nuclear borders and intensity values of non-photoconverted (green) and photoconverted (red) Eos were measured at the timepoints indicated in Figure 2C. Activity from the tp63 transgene was quantified by calculating the green to red fluorescence intensity ratio within individual nuclei of basal and migratory cells. Data visualization was performed in R using the ggplot2 package to generate a dot plot overlaid with lines connecting the mean tp63 expression values of basal epithelial and migratory cells across the tracked timepoints.

Image tracking was performed using the TrackMate plugin (Ershov et al., 2022; Tinevez et al., 2017) in ImageJ/Fiji (Schindelin et al., 2012). Statistical comparisons and significance values in Figure 5E were calculated using non-parametric Welch’s t-tests on GraphPad Prism Version 10.6.1. Representative trajectories in .xml format from TrackMate were exported to Microsoft Excel, and the DiPer plugin (Gorelik and Gautreau, 2014) was used to plot individual trajectories on XY coordinates (Figure 5D).

Labeled nuclei of basal cell-derived migratory cells in (tp63[BAC]:Gal4FF;UAS:nls-GFP) embryos were time-lapse imaged between 19-31 and 31-43 hpf. Trackmate was used to trace migratory cell trajectories along the trunk, and the Cell Counter plugin was used to quantify the proportion of actively migrating and formerly migrating tp63+ cells. Data visualization was performed in R using the ggplot2 package to generate a dot plot overlaid with lines connecting mean percent values of actively migrating and formerly migrating tp63+ cells at the indicated timepoints in Figure 6A.

## Discussion

Mucus cells and ionocytes serve critical functions in all mucosal epithelia (Ma et al., 2018; Shah et al., 2022). In most organs, these specialized cell types are broadly dispersed in an epithelium, rarely clustering or even abutting cells of the same type. This broad distribution is presumably important for their functions in mucus production and ion transport, since mucus coverage and ionic balance must be evenly distributed to maintain tissue health. How these cells transit into the superficial layer has not been well characterized in any organ. We found that in the developing zebrafish epidermis, migratory mucus cell and ionocyte precursors emerge primarily from the ventral embryo and disperse by migration through the epithelium. Our findings in the zebrafish skin suggest that a similar process of emergence from basal cells, cell migration, mutual repulsion, and apical intercalation may underlie the distribution of these cells in other epithelial organs.

The zebrafish epidermis provides a model for observing the entire “life cycle” of developing mucus cells and ionocytes in live animals, from the onset of migration through intercalation. The emergence of migratory cells from basal cells resembles EMT, but, notably, mucus cell and ionocyte precursors begin migrating when epidermal basal cells in zebrafish are just starting to acquire mature epithelial characteristics, including a basement membrane and polarized junctions (Cokus et al., 2019; Le Guellec et al., 2004; O’Brien et al., 2012). At least one feature of mucus cell and ionocyte emergence has not commonly been described in EMT–the coupling of cell division to the onset of migration. Cell division is associated with cell dispersal in the kidney, but in that case cells reinsert immediately after division, thus dispersing only short distances (Packard et al., 2013). Although not commonly associated with EMT, coupling cell division with the onset of migration may be advantageous, since both processes require cells to reduce adhesion with surrounding cells.

By contrast to the immature state of the basal epithelium at the beginning of migration, the periderm has mature apico-basal polarity, adherens junctions, desmosomes and tight junctions (Cokus et al., 2019; O’Brien et al., 2012) when mucus cell and ionocyte precursors intercalate. Intercalation thus requires cells to wedge themselves into a mature epithelium, a process that closely resembles intercalation by multiciliated cells in the developing xenopus epidermis, which involves mutual inhibition (Chuyen et al., 2021), probing between epithelial cells with filopodial protrusions (Ventura et al., 2022), pushing apart epithelial cells with an expanding apical actin ring (Sedzinski et al., 2016), and affinity for epithelial adhesion molecules (Chuyen et al., 2021; Ventura et al., 2022). These similarities suggest that common mechanisms regulate the intercalation of multiciliated cells, mucus cells, and ionocytes. The diversity of mucus cell and ionocyte subtypes in the epidermis raises questions about how multiple cell types coordinate and compete for space in an epithelium.

The zebrafish epidermis is also an ideal setting in which to study mechanisms of cell motility within epithelia. Mesenchymal cell motility on ECM substrates has been well studied (SenGupta et al., 2021), and several studies have investigated mechanisms of trans-epithelial migration by immune cells (Friedl and Weigelin, 2008), but relatively few have focused on mechanisms of migration entirely within epithelia, despite the fact that metastatic cancers can migrate in intraepithelial environments (Eckert et al., 2016; Lee et al., 2021; Schneider et al., 2017; Simon et al., 2001). Several features of mucus cell and ionocyte precursors suggest that they use pseudopod-based amoeboid motility to move through the epithelium, including their presence in a low-ECM environment and their use of leading edge actin-containing protrusions (SenGupta et al., 2021; Welch, 2015). Their migration speeds are relatively slow compared to blebbing amoeboid cells, but similar to some amoeboid cells migrating with pseudopods (Liu et al., 2015; Welch, 2015).

The developing epidermis is increasingly appreciated to contain several types of migratory cells–including sensory axons (O’Brien et al., 2012; Rosa et al., 2023), melanoblasts (Richards et al., 2026), Merkel cell precursors (Craig et al., 2025), T-cells (Robertson et al., 2025), and neutrophils (Schrope et al., 2026). Although all of these cells migrate through developing epidermis, they have different migratory characteristics. Melanoblasts in mice, like mucus cell and ionocyte precursors in zebrafish, migrate above basal cells, but they frequently move back and forth between the intraepithelial space on the apical side of basal cells and the basement membrane-contacting space below basal cells (Richards et al., 2026). Melanoblasts have actin-based protrusions, but elongated, dendritic morphologies that suggest they use a different motility mechanism (Richards et al., 2026). Neutrophils in the developing zebrafish epidermis also have dendritic morphologies and migrate primarily between lateral membranes of basal cells, where they can contact the ECM (Schrope et al., 2026). T-cells in zebrafish migrate in the same territory above basal cells as mucus cell and ionocyte precursors, but appear about a week later in development, and migrate using a stable bleb motility mechanism, with actin concentrated in the rear of cells (Robertson et al., 2025). Comparing the motility mechanisms between all of these cell types migrating in the epidermis may help parse the relative influence of intrinsic and extrinsic factors to motility strategies.

Cell migration coupled with contact inhibition of locomotion is a common mechanism to disperse cells through a tissue (Stramer and Mayor, 2017). The distribution patterns of mucus cells and ionocytes, however, cannot solely be explained as a consequence of undirected migration and contact inhibition, since different subtypes of mucus cells and ionocytes concentrate in different parts of the animal: HRCs are confined to the epidermis above the yolk, NARCs are concentrated in the trunk, and mucus cells are most abundant in the head and distal tail (Figure 1B-B”). Ionocyte and mucus cell identities are not determined by local factors at the end of their migration, since ionocytes and mucus cell precursors are specified hours earlier, before migration. The differences in distribution pattern of each subtype could be achieved by several other mechanisms: cell type-specific chemoattraction (or repulsion) to regional cues, different motility properties for each subtype (e.g., slow moving subtypes stay closer to their origin than fast moving subtypes), subtype specific timing of migration and/or integration (e.g., later migrating cells and earlier integrating cells stay closer to their origin), or asymmetric contact inhibition of locomotion (e.g., one cell type repels another, but not vice versa). Determining which of these mechanisms contribute to their distinct distribution patterns may require creating transgenic reporters for mucus cell and ionocyte subtype precursors that are visible at earlier time-points, before migration begins.

Our results describe a mechanism for the initial distribution of mucus cells and ionocytes, but as animals grow, new mucus and ionocyte precursors must be generated from stem cells, distributed through the expanding epidermis, and remodeled to accommodate developmental transitions in the skin. At juvenile stages (∼3 weeks post-fertilization), the pattern of ionocytes dramatically changes–ionocytes are lost in the skin and concentrate in the gills (Hwang and Chou, 2013; Kowalewski et al., 2021). Also at juvenile stages, as part of a metamorphic transition of the epidermis, the entire periderm peels off (Guzman et al., 2013), presumably bringing along with it embedded mucus cells and ionocytes, which must be replaced. Changing the ionic environment can promote homeostatic changes in the number of ionocytes, also requiring additional proliferation and distribution of new cells (Li et al., 2021; Liu et al., 2018; Peloggia et al., 2021; Peloggia et al., 2024; Xin et al., 2019; Xin et al., 2021). All of the processes described in this study during embryonic stages–emergence from basal cells, intra-epithelial migration, differentiation, and intercalation, likely contribute to these remodeling events. Understanding how mucus cell and ionocyte dispersal is regulated during each of these plastic processes could help us understand the cellular reorganization that must occur in damaged mucosal organs during tissue repair.

---

**Supplementary Figure 1.**
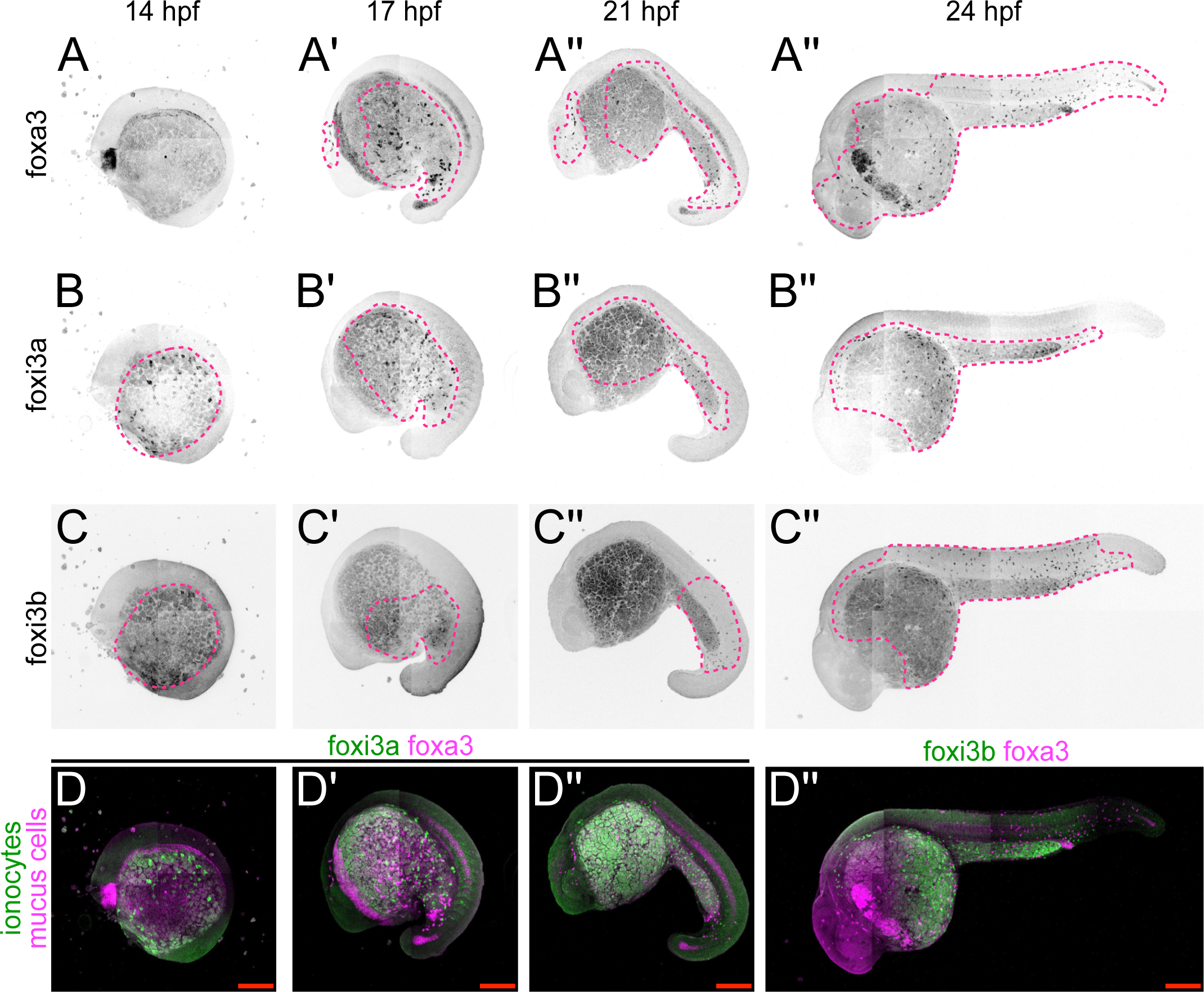
Expression time-course of ionocyte and mucus cell specification genes. RNA-scope in situ hybridization for mucus cell (foxa3) and ionocyte (foxi3a, foxi3b) specification genes at the indicated time-points (A-C’’). Pink dotted lines indicate rough areas containing cells expressing the indicated marker. D-D’’) Merge of foxa3 and foxi3a or foxi3b, as indicated. Foxi3a is shown at earlier stages and foxi3b at 24 hpf since foxi3a fades and foxi3b expression increases over time. Images in D-D’’ are also shown in Figure 1. Scale bars in merged images = 100 μm.

**Supplementary Figure 2.**
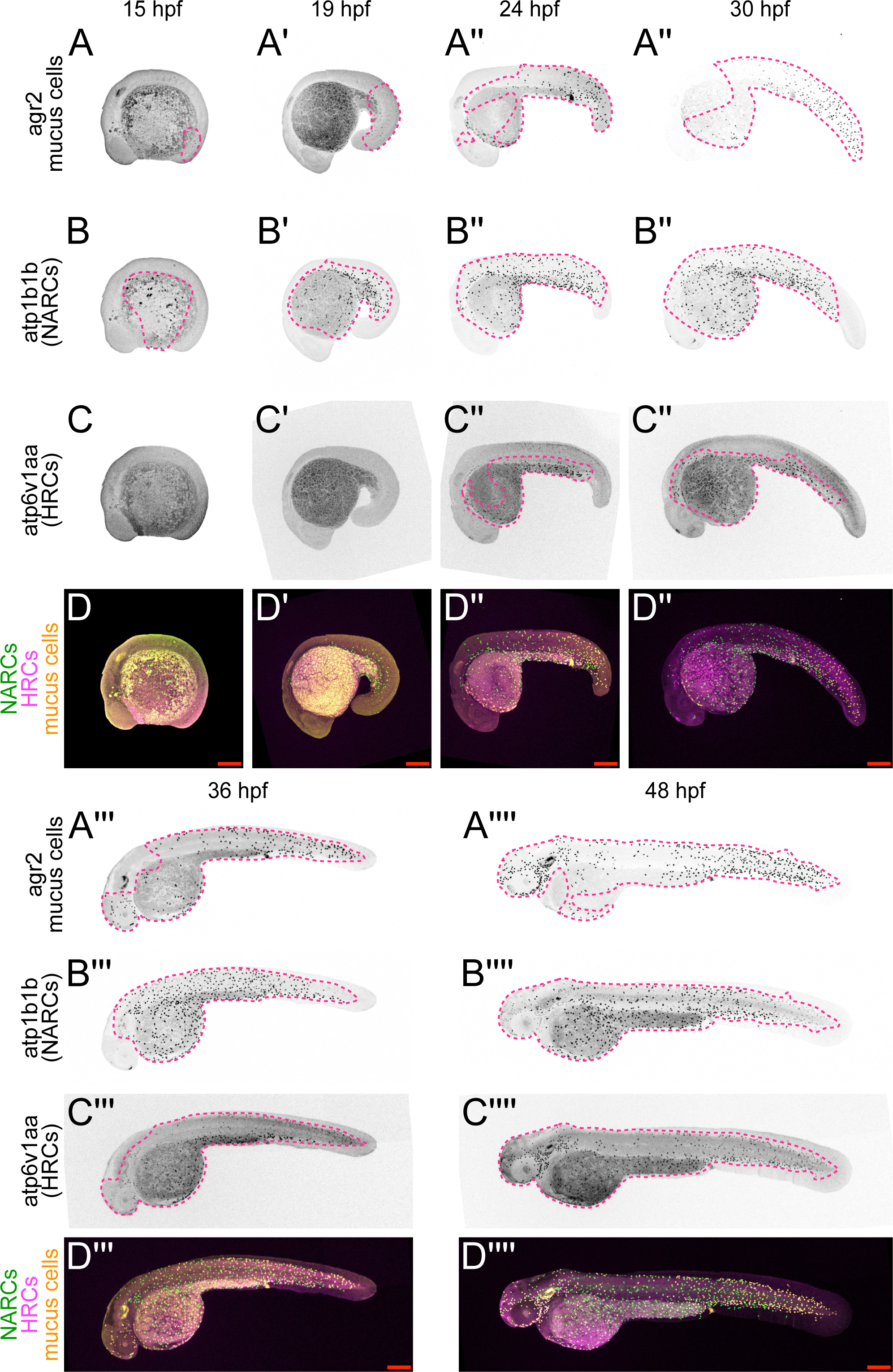
Expression time-course of ionocyte and mucus cell differentiation genes. RNA-scope in situ hybridization for mucus cell (agr2) and ionocyte (atp1b1b, atp6v1aa) differentiation genes at indicated time-points (A-D’’’’). Pink dotted lines roughly show areas containing cells expressing the indicated marker. D-D’’’’) Merge of agr2-expressing mucus cells, atp1b1b-expressing NARCs, and atp6v1aa-expressing HRCs. Note that since all three reporters are expressed in mucus cells at 48 hpf, each cell type can only be distinguished in the merge (D’’’’), where ionocytes are identifiable by their lack of agr2 expression. 48 hpf images (A’’’’-D’’’’) are also shown in Figure 1. Scale bars in merged images = 100 μm.

**Supplementary Figure 3.**
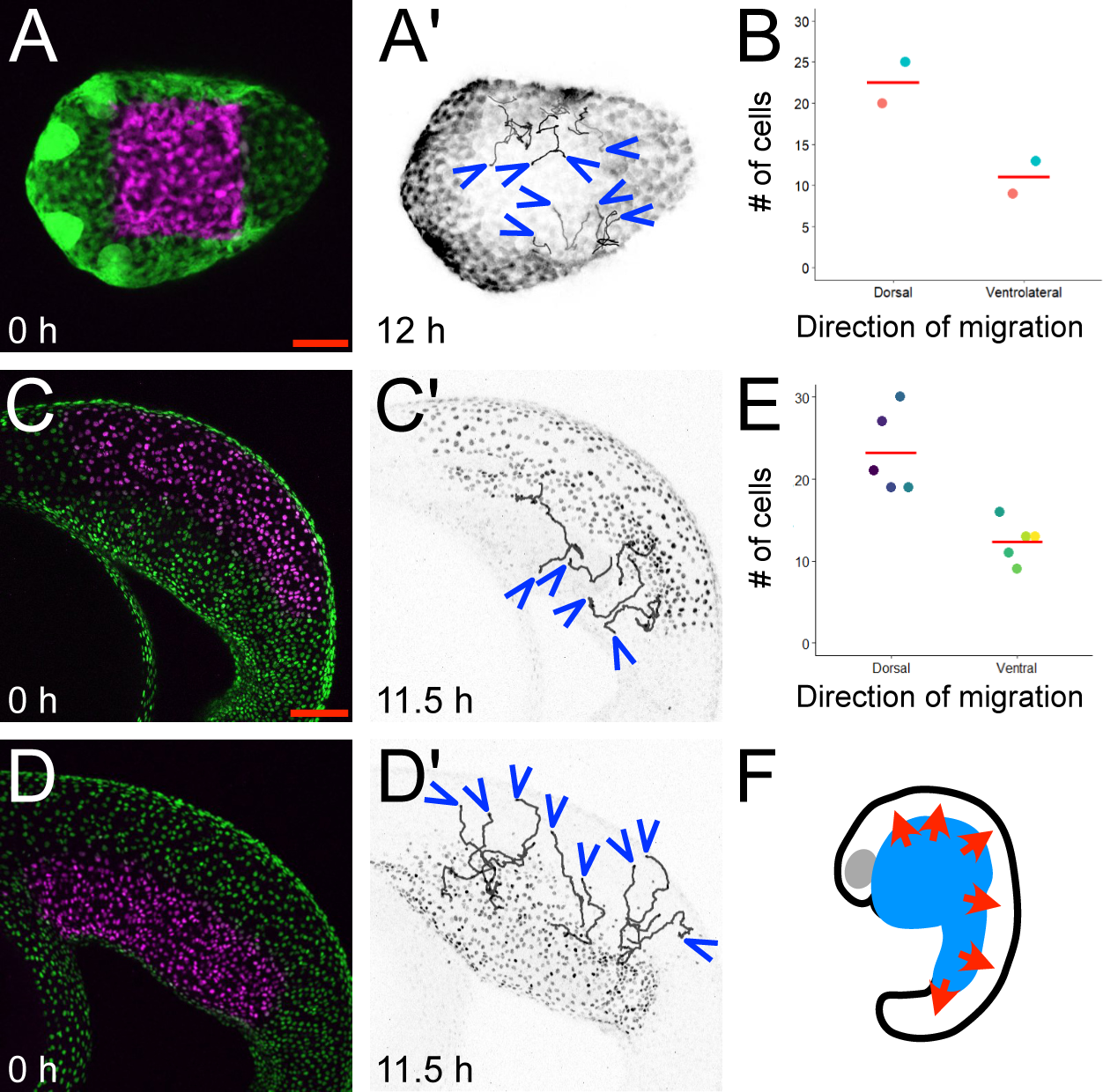
Photoconversion maps migratory cell trajectories. A-A’) A region of the head was photoconverted in animals expressing a photoconvertible fluorescent protein in basal and migratory cells (tp63[BAC]:Gal4FF; UAS:KikGR) at 24 hpf and imaged for 12 hours. A shows photoconverted (magenta) and unphotoconverted (green) cells at the start of the movie; A’ shows the non-photoconverted channel with tracings of individual cell trajectories at the final time-point. Arrowheads indicate the position of cells at the end of the movie. B) Quantification of cells moving into (ventrolateral-to-dorsal) and out of (dorsal-to-ventrolateral) the photoconverted region in the head. Colored dots represent the number of cells moving in each direction from two movies. Dots with the same colors are from the same movie. C-E) Either the dorsal trunk (C-C’) or ventral trunk (D-D’) was photoconverted in animals expressing a photoconvertible fluorescent protein in basal and migratory cell nuclei (tp63[BAC]:Gal4FF; UAS:nls-Eos) at 20 hpf and imaged for 11.5 hours. C and D shows the photoconverted (magenta) and unphotoconverted (green) cells at the start of the movie; C’ and D’’ show just the photoconverted channel with tracings of individual cell trajectories at the final time-point. Arrowheads indicate the position of cells at the end of the movie. E) Quantifications of cells moving out of the photoconverted region in the trunk. Only photoconverted cells were counted in 5 movies each for dorsal and ventral photoconversion. Colored dots represent the number of cells moving in the indicated direction, but, unlike in B, separate movies were used for dorsal and ventral photoconversion experiments. F) Model: cells spread predominantly dorsally away from the ventral embryo.

## Movie Legends

**Movie 1. Photoconverted cells migrate away from the yolk extension.** Basal cells in the yolk extension region were photoconverted at ∼18 hpf and time-lapsed starting at ∼20 hpf. Individual photoconverted cells (magenta) can be seen migrating dorsally away from the yolk extension. Cell trajectories from this movie are summarized in Figures 1J. Time-stamp is in the upper right. Scale bar = 100 μm.

**Movie 2. Migratory cells emerge from tp63-expressing basal cells following cell division.** Two examples of migratory cells emerging from basal cells after cell division. Movies correspond to Figures 2A and B. Cell paths are traced in green. Time-stamp is in the upper right. Scale bar = 20 μm.

**Movie 3. Cells expressing a vimentin reporter migrate away from the yolk extension.** Cells expressing a vimentin reporter. Movies correspond to Figures 2E and F. Representative paths of migratory cells are traced. Time-stamp is in the upper left. Scale bar = 50 μm.

**Movie 4. Migrating basal cell-derived cells begin expressing ionocyte and mucus cell reporters during migration.** Basal cell-derived migratory cells co-expressing reporters for HRC ionocytes or mucus cells during migration. Movies correspond to Figure 3A and B. Time-stamp is in the upper right. Scale bar = 20 μm.

**Movie 5. Cell morphology during migration.** Two examples of migrating mucus cell precursors. Movies correspond to Figures G-H. Time-stamp is in the upper right. Scale bar = 10 μm.

**Movie 6. Migrating cells repel one another.** Migrating mucus cell precursors change direction after contacting one another. Arrows show trajectories of each cell; stars show instances of contact. Scale bar = 10 μm.

**Movie 7. Migrating cells make actin-based protrusions at their leading edges.** Migrating cells mosaically labeled with LifeAct-GFP. Time-stamp is in the upper right. Scale bar = 10 μm.

**Movie 8. Migratory cells proliferate as they migrate.** Part 1: Basal cell-derived migratory cells expressing a nuclear (histone) marker pause migration to divide. Part 2: Mucus cell precursors pause migration to divide. Time-stamps are in the upper right. Scale bars = 10 μm.

**Movie 9. Migratory cells intercalate into the periderm epithelium.** Migratory cells mosaically expressing an actin reporter as they intercalate into the periderm. Movie corresponds to Figure 3A and B. Time-stamp is in the upper right. Scale bar = 20 μm.

## Acknowledgments

We thank Anna Huttenlocher (University of Wisconsin), Sheng-Ping Hwang (Academic Sinica), Gage Crump (University of Southern California), and Laura Yatime (University of Montpelier) for transgenic lines (See Table 1 in Materials and Methods). We are particularly grateful to Jieun Park and Michel Bagnat for sharing the Occludin BAC reporter line before publication. We thank staff from the following UCLA core facilities for advice and help with imaging, cell sorting and sequencing, respectively: UCLA Broad Stem Cell Research Center Microscopy Core (RRID:SCR_024914), UCLA Broad Stem Cell Research Center Flow Cytometry Core, UCLA Technology Center for Genomics & Bioinformatics (TCGB). We thank Linda Dong and the UCLA zebrafish core facility staff for expert animal care. This work was supported by a National Institute of Arthritis and Musculoskeletal and Skin Diseases (NIAMS) grant (R01AR064582) to AS and a NIAMS post-doctoral fellowship (F32AR086666) to OJ.

